# Standardised empirical dispersal kernels emphasise the pervasiveness of long-distance dispersal in European birds

**DOI:** 10.1101/2021.11.08.467775

**Authors:** Guillermo Fandos, Matthew Talluto, Wolfgang Fiedler, Robert A. Robinson, Kasper Thorup, Damaris Zurell

## Abstract

1. Dispersal is a key life-history trait for most species and essential to ensure connectivity and gene flow between populations and facilitate population viability in variable environments. Despite the increasing importance of range shifts due to global change, dispersal has proved difficult to quantify, limiting empirical understanding of this phenotypic trait and wider synthesis.
2. Here we aim to estimate and compare empirical dispersal kernels for European breeding birds considering average dispersal, natal (before first breeding) and breeding dispersal (between subsequent breeding attempts), and test whether different dispersal properties are phylogenetically conserved.
3. We standardised and analysed data from an extensive volunteer-based bird ring-recoveries database in Europe (EURING) by accounting for biases related to different censoring thresholds in reporting between countries and to migratory movements. Then, we fitted four widely used probability density functions in a Bayesian framework to compare and provide the best statistical descriptions of the average, the natal and the breeding dispersal kernels for each bird species.
4. The dispersal movements of the 234 European bird species analysed were statistically best explained by heavy-tailed kernels, meaning that while most individuals disperse over short distances, long-distance dispersal is a feature in almost all bird species. The overall phylogenetic signal in both median and long dispersal distances was low (Pagel’s λ < 0.40), implying a high degree of taxonomic generality in our findings. As expected in birds, natal dispersal was 5 Km greater as an average than breeding dispersal for most species (88% species analysed).
5. Our comprehensive analysis of empirical kernels indicates that long-distance dispersal is common among European breeding bird species and across life stages. The dispersal estimates offer a first guide to selecting appropriate dispersal kernels in range expansion studies and provide new avenues to improve our understanding of the mechanisms and rules underlying dispersal events.

## Introduction

Animal dispersal describes the movement from birth to breeding patch (natal dispersal) or between successive breeding patches (breeding dispersal) and is a fundamental biological process in ecology and evolution (Greenwood, 1980). Dispersal is a crucial determinant for different ecological processes at a wide range of spatial and temporal scales. At a macro scale, dispersal plays a key role in determining evolutionary patterns of speciation and extinction and the geographical distribution of species (Bowler & Benton, 2005; Kisel & Barraclough, 2010). Within populations, dispersal plays a key role in the genetic structure of populations and meta-population dynamics through its direct contribution to gene flow (Bonte & Dahirel, 2017; Hallatschek & Fisher, 2014; Venail et al., 2008) and in maintaining local populations (Millon et al., 2019; Schaub & Ullrich, 2021). Improved understanding of dispersal across many species is becoming increasingly important, given the need to predict how populations will respond to global change (Barbet□Massin et al., 2012; Zurell, 2017). Despite this broad relevance, however, we still have a limited understanding of this phenotypic trait as standardised empirical data on animal dispersal are largely missing, hampering wider synthesis of mechanisms and underlying drivers (Bullock et al., 2017).

Quantifying how far and how often animals move across the landscape is extremely challenging (Nathan, 2001). More recently, understanding of movement processes has advanced through the implementation of new molecular tools (Hobson, 2005; Woltmann et al., 2012) and the use of cutting-edge biotelemetry (Kays et al., 2020; Kranstauber et al., 2011). Still, empirical dispersal measurements on vertebrates are scarce, mostly constrained to few organisms, and geographically limited (Paradis et al., 1998). As a consequence of these challenges, comparative dispersal analyses across species have relied on standardised biometric indices as proxies to quantify dispersal ability (Dawideit et al., 2009; Sheard et al., 2020), or imputation methods that fill information gaps based on phylogenetic relatedness between species (Barbet□Massin et al., 2012).

Syntheses of field movement and dispersal data provide a promising avenue for overcoming empirical data limitations for many vertebrate species and large spatial extents (Tucker et al., 2018). For example, two decades ago, Paradis et al. (1998) estimated average natal and breeding dispersal distances for 75 British bird species based on nearly 100 years of ringing data. This analysis explored how dispersal distances vary according to certain life-history traits (e.g. migratory behaviour, range size, habitat) and dispersal type (breeding or natal dispersal). These estimates have subsequently been used to project bird dispersal and range dynamics under climate change (Barbet□Massin et al., 2012). However, the original dispersal estimates by Paradis et al. (1998) were constrained to Great Britain, to only a subset of European breeding birds, and summarised only average dispersal distances rather than explicitly estimating dispersal kernels and analysing their shapes. Dispersal kernels, which represent the density of dispersing individuals at certain distances from the source, provide a better understanding of the mechanisms and rules underlying dispersal events and are a prerequisite for modelling spatial population dynamics for scenarios of global change (Bullock et al., 2017; Nathan et al., 2012; Paradis et al., 2002). Yet, building a large dataset of empirical dispersal kernels for a wide range of species in large areas is challenging due to different biases and uncertainties in the field observations (Nathan et al., 2012).

Different studies have implemented a variety of functions to represent the frequency distribution of the dispersal distances (Exponential, Nathan et al., 2012; Gamma, van Houtan et al., 2007; Half-Cauchy distribution, Paradis et al., 2002; Weibull, Nathan et al., 2012). These functions differ in the shape of the dispersal kernel and thus in the relative probability of different dispersal distances with consequent implications for predicting range change. Functions like the exponential kernel are popular as they have an underlying theoretical basis that represents movement in a random direction with a time or distance-dependent settlement rate (Bullock et al., 2017; Nathan et al., 2012). By contrast, heavy-tailed kernels such as the Half-Cauchy, Gamma and Weibull distribution assume a combination of local and distant selective pressures and they expect that a few individuals fly long distances (Viswanathan et al., 1996). To date, only a few studies compared different dispersal kernel functions on birds (Nathan et al., 2012; Paradis et al., 2002; Van Houtan et al., 2007, 2010). These indicated that simple summary statistics of empirically measured dispersal distances (rather than estimating dispersal kernels based on probability distributions) underestimate the species’ dispersal ability and that heavy-tailed kernels may best explain empirical dispersal patterns (Paradis et al., 2002; Van Houtan et al., 2007). Comparing the performance of alternative empirical dispersal kernels for large numbers of species will improve our ecological understanding of relevant dispersal processes and their proximate and ultimate causes (Stevens et al., 2014).

Here, we aim to quantify empirical dispersal kernels of breeding birds across Europe, compare the dispersal characteristics of natal and breeding dispersal, and test for phylogenetic signal in different dispersal metrics. We use data on marked birds from EURING – The European Union for Bird Ringing database – that holds several million records of European bird movements (Du Feu et al., 2016). Although a uniquely rich data source on bird movements, analysis of dispersal distance based on EURING data is challenging because dispersing and migrating birds are not separated, and sampling effort is heterogeneous (Paradis et al., 1998; Korner-Nievergelt et al 2010). Therefore, we develop a methodological framework that addresses these potential biases. Based on this, we first estimate dispersal kernel parameters for average dispersal (pooling all age stages when it was not possible to separate them) as well as for breeding and natal dispersal using four different probability density functions and assess the best-fitting one. Then, we calculate multiple descriptors of dispersal (e.g. median and maximum dispersal distances) and quantify the phylogenetic signal in these descriptors. The use of multiple dispersal descriptors will allow us to disentangle different selective pressures on short-versus long-distance dispersal patterns (Claramunt, 2021; Sheard et al., 2020),

## Methods

### Ringing data

Raw data on dispersal distances were obtained from the EURING database (Du Feu et al., 2016). The data were requested following an approach that allowed us to keep only the reliable observations and test for different sampling biases. Therefore, for the present study, we included distances between the ringing and re-encounter locations of birds ringed and subsequently re-encountered between April and July (which represents the core breeding season for most species) from 1979 until 2018 from almost all ringing schemes in Europe (see supplemental material 1). Re-encounters within the same breeding season as ringing were excluded. When multiple subsequent encounters at the same coordinates as the previous encounter were available, only the first one was considered. We re-classified the field codes for the condition of the reencountered birds into two classes, dead (code: 1-3) and alive (code: 4-8), and defined two age classes with respect to the age of the birds when ringed: juvenile for birds ringed in their year of birth (age code 1 and 3), and adult for birds ringed later than the first year of birth.

Because sampling effort varies across schemes and species, we selected a balanced dataset in terms of sample size across Europe for all species, age groups (nestling or adult), and types of recovery (dead or alive) that allowed us to estimate dispersal and tackle the uneven spatial coverage and heterogeneous sampling associated with different types of re-encounter. In particular, we used a stratified random sampling by 5° grid cell to select ringing site locations across Europe, then chose a minimum of 20 records and a maximum of 100 records per 5° grid cell with c. 60% dead recoveries and 40% alive recoveries where possible. Only recoveries where the location of the encounter was known to a precision of ±5km were included. The data were further screened following the procedure described in Paradis et al. (1998) to remove spurious effects and heterogeneity as far as possible (birds in poor condition, ringing or recovered events in uncommon circumstances, and lack of accuracy on the dates and places of ringing and/or recovery). Several species are not separated in sex classes in the database; hence, we avoid to use sex as a category in this study. In total, the ringing data obtained from EURING consisted of 602,703 ringing and re-encounter events from 273 species.

### Potential bias analysis

Ringing databases hold dispersal information that could not be acquired using alternative techniques. Ring-recovery data are available for many species and are not constrained by sampling being restricted to particular locations (Tellería et al., 2012). However, drawing conclusions on dispersal from raw data can be misleading because re-encounters, and hence dispersal distances, are the result of a heterogeneous observation process and subject to strong sample biases (Fandos & Tellería, 2018; Korner-Nievergelt et al., 2010; Naef□Daenzer et al., 2017; Thorup et al., 2014). Here, we used different approaches to exclude data that can lead to potential biases in the calculation of dispersal for the different species. In particular, those biases related to (i) different recovery rates between types of recovery (live recaptures, resightings and dead recoveries), (ii) migratory movements and (iii) the minimum number of cases used to infer robust dispersal estimates:

(i) Although a large variation in ringing and recovery effort could potentially bias the spatial and temporal distributions of ringing data, we expect that the large spatial scale of our study can minimise the biases associated with the heterogeneous recovery rates. Nevertheless, dead and alive recaptures may be affected by different biases related to catching effort by ringers and reporting probability (Paradis et al., 1998). For instance, the spatial distribution of birds recaptured alive is likely to differ from dead recoveries as the former depends on the spatial and temporal efforts in field ornithologist activities (more recoveries at places with active research/ringing stations; Tellería et al., 2014), while the latter are mostly reported by the general public and so are more evenly distributed. Therefore, in an exploratory analysis, we compared the dispersal estimates obtained from using different recovery types. Comparison of the results indicated that both dead and alive recaptures (but excluding live resightings), showed similar dispersal patterns (see supplemental material 2).

(ii) The dispersal analysis of migratory or partial migratory species is particularly challenging because of variation in migration phenology between individuals and populations across Europe (Lehikoinen et al., 2019). Because migratory movements may lead to overestimation of dispersal distances, we aimed to exclude individuals captured or recovered during migration in the late or early breeding season, using a two-step approach. First, we estimated the potential core breeding period for each species and each spatial (5°) grid cell in Europe to account for the breeding time variation across space. For this, we used generalised additive models (GAMs) to regress dispersal distance against a smoothed function of the time of the year and used the second derivative to distinguish peak migratory periods with sudden increases in dispersal distances from the core breeding season with comparably stable dispersal distances. Second, we used the 95% quantile of the distances observed in the core breeding period as a conservative cut-off distance to distinguish between dispersal events and migratory movements (Supplemental material 3).

(iii) Finally, we ran an exploratory analysis, where we used different subsets of ring-recoveries to assess how the number of events would affect the dispersal estimation. We concluded that a minimum of 20 individuals per analysis was sufficient to ensure robust dispersal estimates (ensuring a minimum sample size of *n*=10 per parameter in two-parameter dispersal kernels).

### Statistical modelling of dispersal distance kernels

For each species, we fitted an average dispersal kernel (not distinguishing natal and breeding dispersal), and if enough data were available, we additionally fitted a natal dispersal kernel and a breeding dispersal kernel. We used a Bayesian approach to fit four commonly used dispersal kernel functions in their one-dimensional form (i.e. probability density functions) directly to the distribution of dispersal distances (Table 1). These four 1- or 2-parameter probability density functions have been commonly used in analysing bird dispersal data (Nathan et al., 2012). Overall, we fitted average dispersal kernels for 234 species. Because of sample size issues, natal dispersal kernels and breeding dispersal kernels were fitted only for 113 and 122 species, respectively; thus we estimated 1,876 dispersal kernels for the combinations of species x four dispersal functions x average /natal/breeding dispersal events.

**Table 1.**
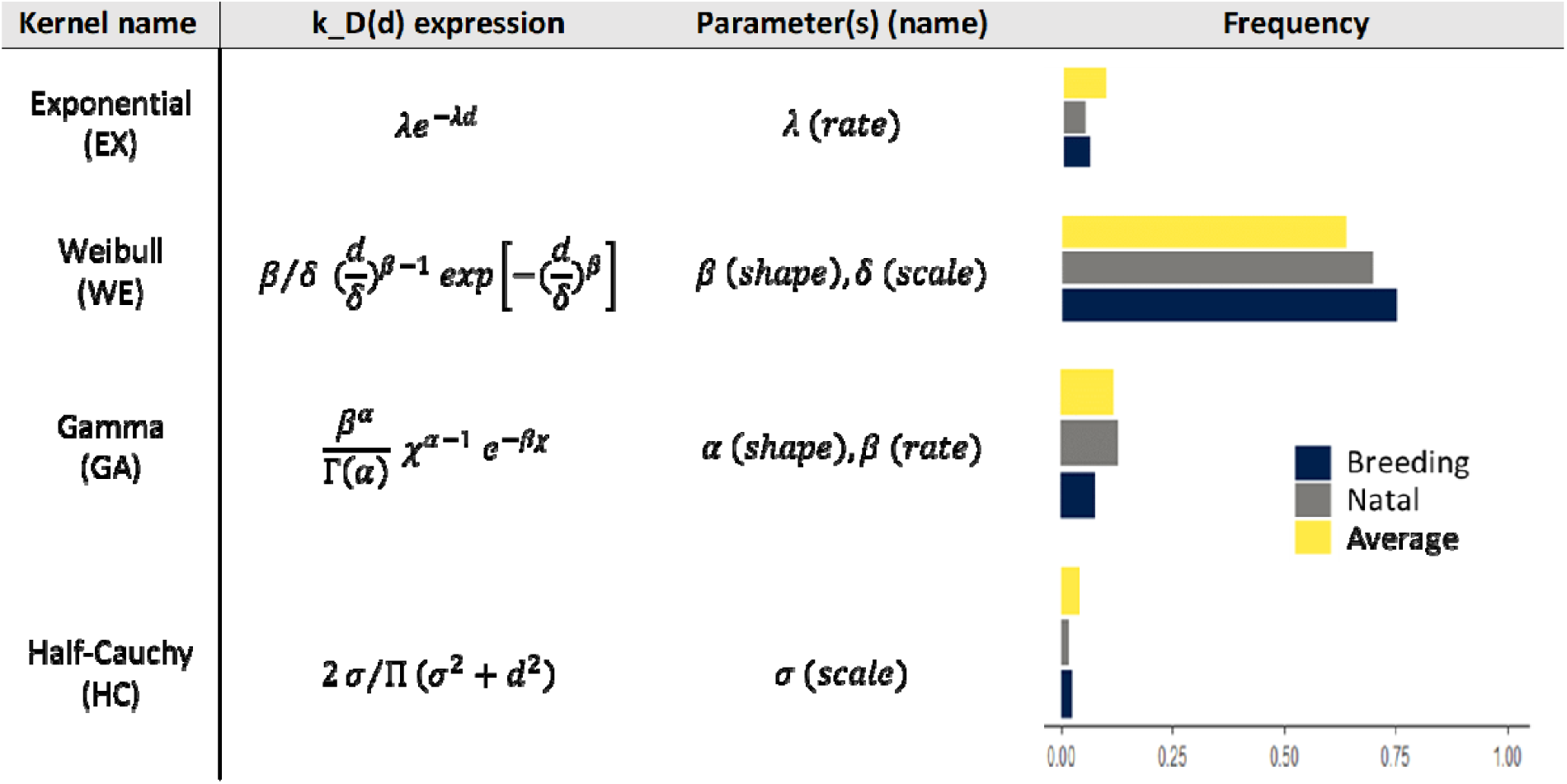
Alternative probability density functions to estimate dispersal kernels k for European birds. We provide the expressions of the one-dimensional dispersal distance kernels k_D_ as function of the distance d, as well as the parameters to estimate for each function. The frequency corresponds to the posterior model probabilities from the computed marginal log-likelihoods via bridge sampling divided by the number of species (frequency = 1 indicates the most likely distribution). The three bars represent the frequency with which each dispersal kernel best fitted the different dispersal types (average : yellow, breeding: blue and natal: grey)

One of the main challenges of fitting dispersal kernels to the EURING database for dispersal analysis is that different schemes have different procedures for reporting birds ringed and subsequently encountered again (Du Feu et al., 2016). For example, some schemes have minimum distances before a bird’s re-encounter will be deemed reportable. This means that recaptures below a specific distance from the ringing location are not always reported, and this lower threshold of reporting a recovery varies between schemes. The resulting bias of omitting short dispersal events is problematic because it affects the dispersal kernel’s shape. For overcoming this problem, we defined two kinds of observation. When the dispersal distance is 0 m, we specified the observation as potentially *censored*. When the observation is precisely known and greater than 0 m, we defined it as *accurate*. Preliminary analyses showed that France had a particularly high threshold for reporting recoveries, but the thresholds for the other schemes also seemed variable. To avoid any arbitrary choices for the censoring thresholds, we decided to infer these from the model.

In the following, we describe the steps to estimate the scheme-specific censoring thresholds and fit the four probability density functions (distributions) to our empirical data (Figure 1; see code *availability*). The procedure was carried out separately for average, breeding and natal dispersal.

**Figure 1:**
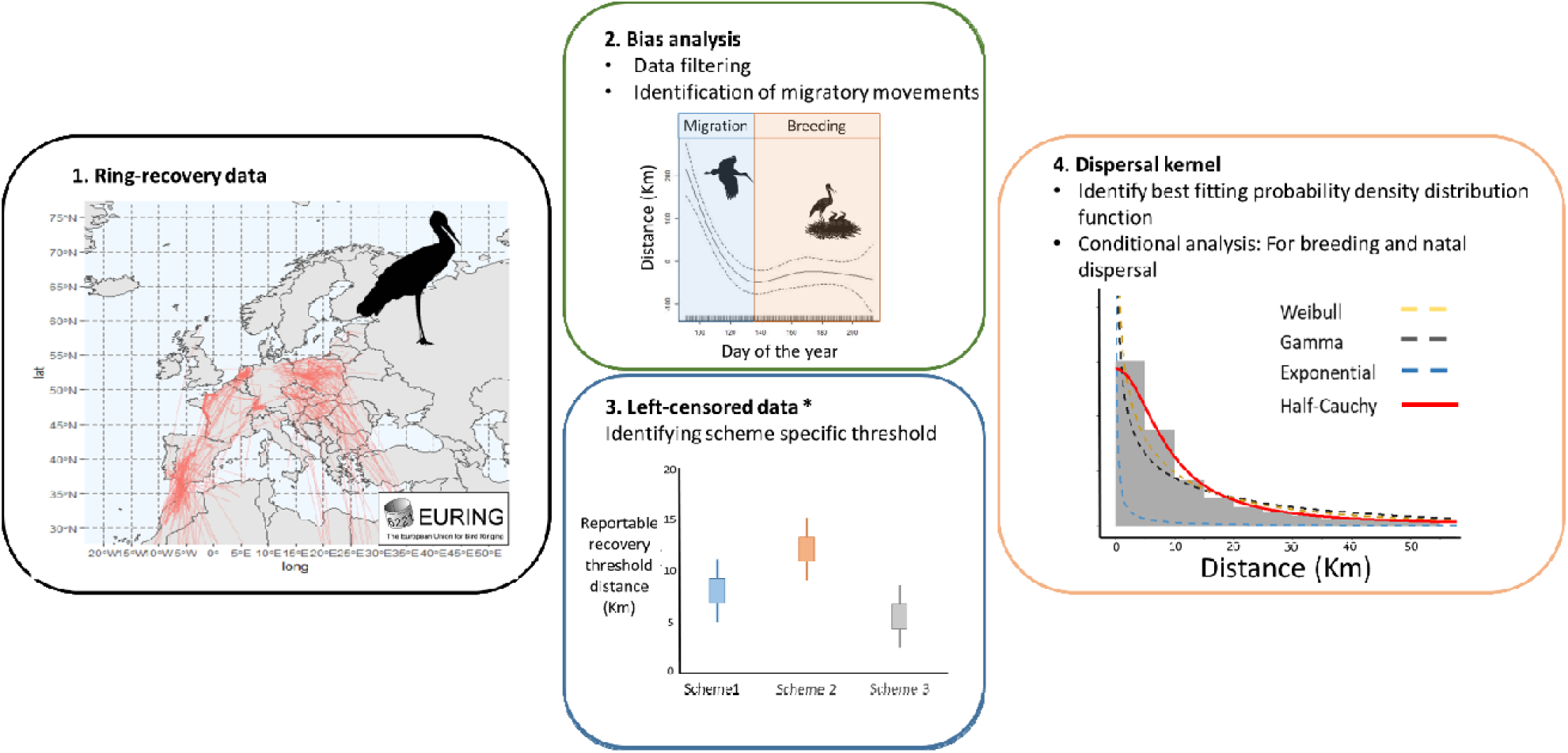
Estimating specific dispersal parameters (White Stork, *Ciconia ciconia* as an example). 1) A spatially balanced data set per species was requested from EURING. 2) Data screening included potential bias analysis accounting for the different recovery rates between recovery types (live recaptures, resightings and dead recoveries), and migratory movements. 3) Scheme-specific thresholds for the reported recovery threshold distance were estimated. Finally, 4) four different density distributions (Exponential, Gamma, Weibull and Half-Cauchy were fitted to all species, and the best fitting distribution was identified for each species.

1. To make use of maximum information for identifying the scheme-specific censoring thresholds, we first fitted a separate dispersal kernel for each specie, with a shared parameter describing the threshold for each scheme. We repeat this process for each dispersal function (Exponential, Gamma, Weibull, Half-Cauchy). We selected the best-fitting distribution by computing the marginal log-likelihood via bridge sampling for each model and computing the posterior probability with the bridgersampler R package (Gronau et al., 2020). Finally, using this best model, we estimated the posterior distribution of the scheme-specific censoring threshold parameter.

2. We used the posterior distribution of the scheme-specific threshold parameter from the previous step as an informative prior in single-species models and for each dispersal function. The objective of these models was to estimate the dispersal kernels for each species, given the degree of left-censoring, compute the posterior model probabilities from marginal likelihoods, and assess which distribution is the “best” for each species using the marginal log-likelihood via bridge sampling. For all species and dispersal functions, a) we extracted the dispersal kernel parameters (the mean and the credible interval of each parameter), b) we derived the empirical median dispersal distance (and the 95% credible interval for the median) analytically from the dispersal kernels, and c) derived long-distance dispersal measures, which we defined as the 95% percentile from a posterior predictive dispersal simulation with the estimated parameters.

### Phylogenetic signal in dispersal estimates

We used multivariate generalised linear mixed models to estimate the phylogenetic dependency in both descriptors of the dispersal ability, the median, and the long-distance dispersal (95% upper percentile of dispersal distances) estimates from the best-fitted distribution for each species. Dispersal estimates were log-transformed to satisfy assumptions of normality and linearity and scaled to have a mean of 0 and a variance of 1. We fitted two separated multivariate Gaussian models for the median and the long-distance dispersal and included phylogenetic relatedness as a random effect. We fitted both models, including no fixed effects and estimated the amount of variation in the dispersal estimates explained by shared ancestry between species (i.e. phylogenetic signal) by calculating the parameter λ (Pagel’s λ; Pagel, 1999).

We also explored the relationship between median versus long-distance dispersal by fitting multivariate generalised linear mixed models, with the median dispersal distance as a response variable, the long-distance as a fixed effect and the phylogenetic relatedness as a random effect. All models were implemented in a Bayesian framework using Markov chain Monte Carlo (MCMC) sampling in the package MCMCglmm (Hadfield, 2010) in R version 4.0.5. We ran all models with three chains and 100 000 iterations, with a burn-in period of 1000 and a sampling interval of 50. The convergence of the models was confirmed by examining the effective sample size (greater than 1000) and autocorrelation between samples (less than 0.10) for each chain, as well as the Gelman–Rubin statistics (less than 1.1) among chains. Priors were initially set using inverse-Wishart priors for the phylogenetic and residual variance (V□ =□1, v = 0.002). Parameter estimates from models are reported as the posterior modes with 95% lower and upper credible intervals (CIs). All phylogenetic analyses were conducted on a sample of 100 trees obtained from the Hackett backbone of the global bird phylogeny (www.birdtree.org; Jetz et al., 2012).

### Association between natal and breeding dispersal

We explored the association between natal and breeding dispersal estimates from the best-fitting distributions for each species while accounting for the non-independence of species related to their joint evolutionary history by using a multivariate generalised linear mixed model. We fitted the model using the median natal dispersal distance as a response variable, the median breeding dispersal distance as a fixed effect and phylogeny as a random effect (see above for details about priors and model fitting). We fitted the model for the subset of 108 species where all measures were available. Dispersal estimates were log-transformed to satisfy assumptions of normality and linearity and scaled to have a mean of 0 and a variance of 1.

We ran the same models to estimate the association between the mean dispersal distances reported in Paradis et al. (1998) and our median dispersal estimates from the best-fitting distribution for the subset of 75 species where both measures were available.

## Results

We analysed a total of 563,276 capture-recapture events from 234 species (median capture-recapture event per species n = 419, max = 27’837, min= 21), covering 55 bird families. The four probability density functions converged for all species. The Weibull distribution was the best-fitting function for 156 out of 234 species (Fig. 2; Table 1). The Gamma distribution was the best one for 34 species, the exponential for 32 and the Half-Cauchy for 12 species. We analysed a total of 122 species for natal dispersal, and the Weibull was the best-fitting function for the majority of the species (92 out of 122 species). In the case of the breeding dispersal, the Weibull was the best-fitting function for 88 out of 113 species analysed.

**Figure 2:**
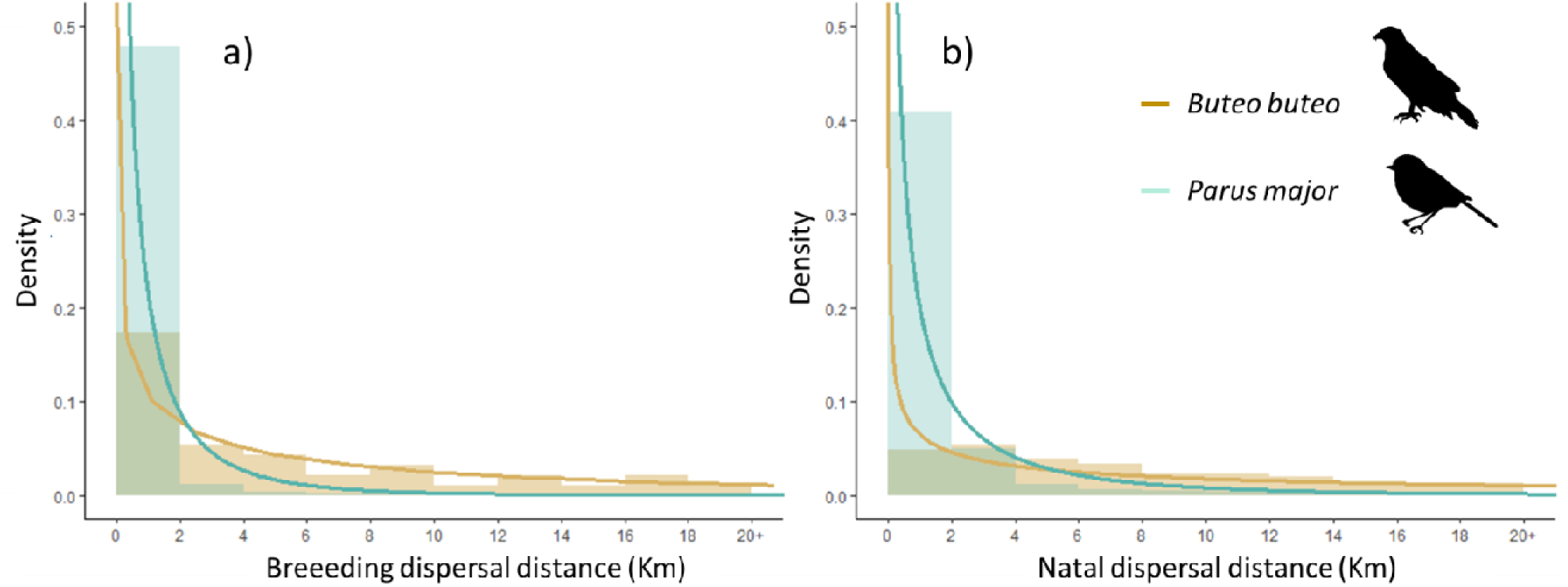
Breeding (a) and natal (b) dispersal kernels for two species: *Parus major* and *Buteo buteo*. Bars represent observed frequency distributions and lines the Weibull probability density curves, which was the best-fitting one for both species.

The dispersal estimates (median and long-distance dispersal) varied between species and species orders (Fig 3; Fig S7.1). The phylogenetic signal for the median dispersal distances was λ = 0.373 [0.115-0.636], whereas the phylogenetic signal for the long-distance dispersal was λ = 0.236 [0.056-0.462]. Reassuringly, the subset of species with large enough sample sizes to estimate breeding (n=122) and natal dispersal (n=113) reflected well the range of dispersal distances found over all species (n=234; Fig. S8.1).

**Figure 3.**
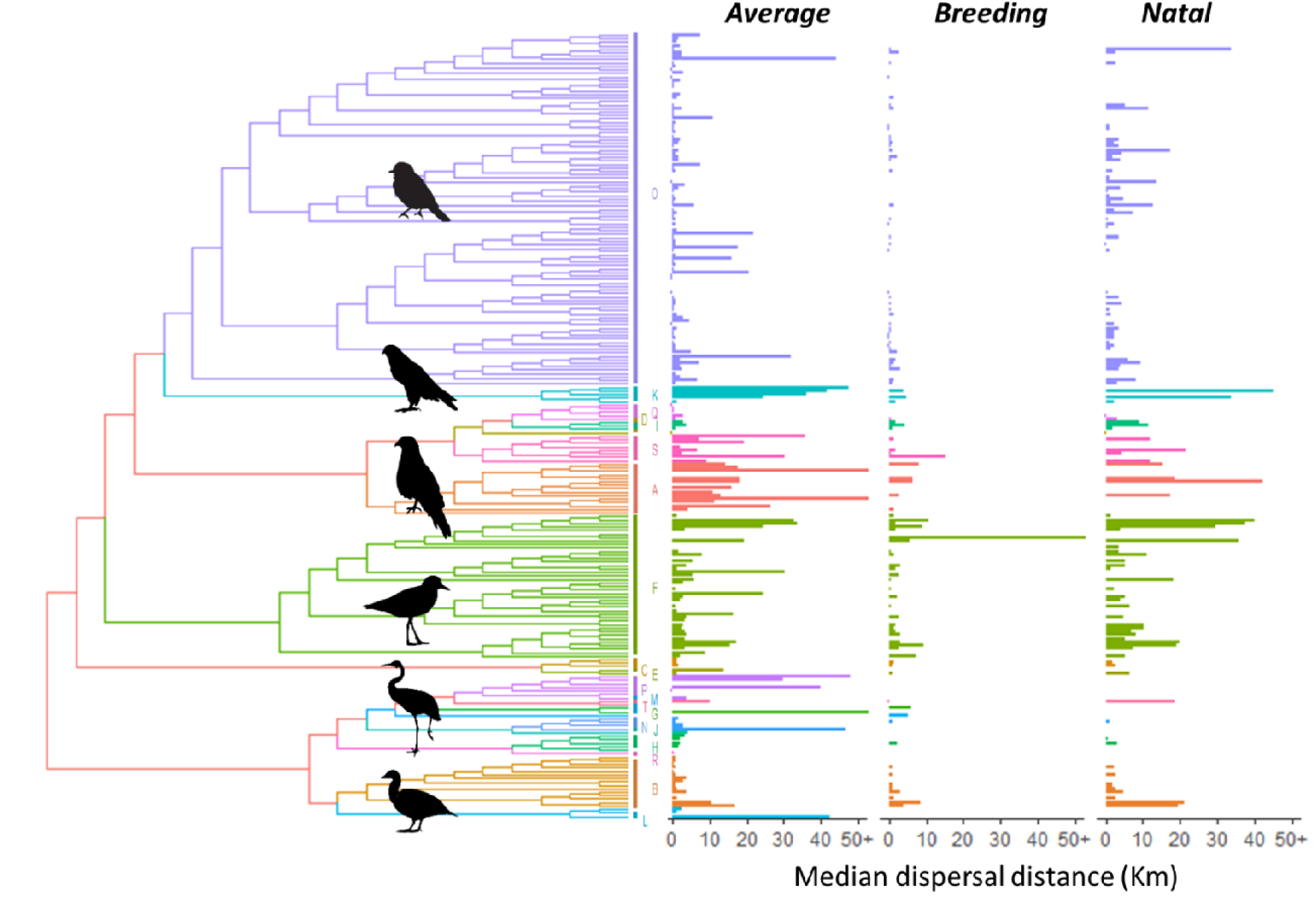
Median dispersal distance (km) from the best-fitting distribution along the bird phylogeny for the average (234 species), breeding (113 species) and natal dispersal (121 species). The dispersal distance is truncated at 50 Km for visualisation purposes. Each colour and letter represent the same Order in the phylogeny and the bar plots. *A: Accipitiformes, B: Anseriformes, C: Apodiformes, D: Bucerotiformes, E: Caprimulgiformes, F: Charadriiformes, G: Ciconiiformes, H: Columbiformes, I: Coraciiformes, J: Cuculiformes, K: Falconiformes, L: Galliformes, M: Gaviformes, N: Gruiformes; O: Passeriformes, P: Pelecaniformes, Q: Piciformes, R: Podicipediformes, S: Strigiformes, T: Suliformes*.

On average, median natal dispersal distances were larger than median breeding dispersal distances (Fig. 4a). Natal and breeding dispersal estimates from the best-fitting kernels had a positive correlation 0.237 (95% CI: 0.036-0.473; pMCMC= 0.039; Fig. 4b). Better correlations resulted when we compared natal and breeding dispersal estimates for the same distribution functions (see figure S5.1 for Weibull distribution). Median dispersal estimates (from the best-fitting kernels) were also significantly correlated with mean dispersal distances reported for n=75 species in Paradis et al. (1998), although the dispersal distances from Paradis et al. (1998) based on summary statistics were larger than our kernel-based estimates (Fig S6.1).

**Figure 4:**
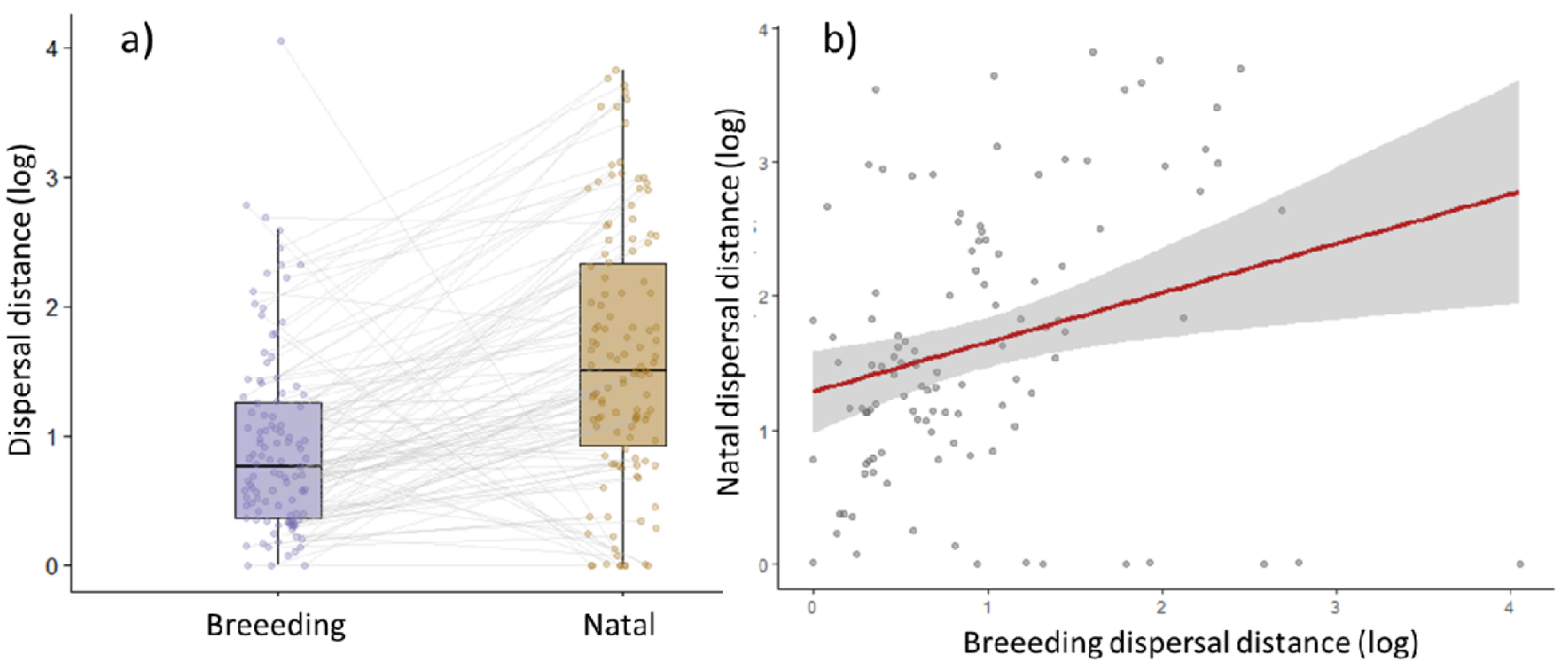
a) Boxplot diagram for the log median dispersal distance (km) from the best-fitting distribution for natal and breeding dispersal. Lines connect the same species in both types of dispersal. c) Linear relationship between breeding and natal dispersal (log).

## Discussion

While much theory has been developed around bird dispersal and their impacts on populations, few empirical studies have estimated and synthesised dispersal kernels for multiple species, a prerequisite for modelling species spatial dynamics (but see Paradis et al., 2002; Van Houtan et al., 2007). In this paper, we estimated average dispersal kernels for 234 bird species across Europe and natal and breeding dispersal kernels for a subset of 122 and 113 species, respectively. This extensive analysis allows an improved understanding of interspecific variations in dispersal patterns and strategies in European birds. Specifically, we found that the dispersal of almost all bird species and across age (natal and breeding dispersal) follows a heavy-tailed distribution, indicating a general tendency towards long-distance dispersal in birds. This result supports previous findings that although most individuals from the different species do not move far, a small proportion of individuals disperse very long distances (Paradis et al., 2002; Van Houtan et al., 2007). More importantly, the phylogenetic signal in dispersal characteristics was weak, indicating that phylogenetic relatedness is a poor predictor of dispersal across bird species but that other internal and external factors may play important roles in determining this phenotypic trait.

Long-distance dispersal events are extremely relevant for population dynamics and range colonisation across changing landscapes, but their low frequency and detectability make them hard to measure and quantify (Clobert et al., 2012; Travis et al., 2013). Empirical dispersal kernels are a fundamental tool to address many of the limitations for characterising dispersal patterns (Bullock et al., 2017; Nathan et al., 2012), in particular when direct measures of dispersal fail to capture the frequency of potential long-distance dispersal events (Koenig et al., 1996; Whitmee & Orme, 2013). The standardisation of dispersal kernels across a wide range of species should allow more realistic and representative forecasts of potential species distributions and better integration of dispersal in comparative life-history analysis (Nathan et al., 2012; Stevens et al., 2012,Bullock et al., 2017). The heavy-tailed distributions probably result from the interplay or overlap of multiple movement modes that widen dispersal kernels when considered simultaneously (Nathan, 2008). Dispersers may switch between movement modes based on the complex trade-offs between internal state, environmental context, motion capacity, and navigational ability (Nathan, 2008). Future analyses will benefit from integrating detailed movement behaviour with improved analytical methods to understand how environmental context affects dispersal, and consequently, eco-evolutionary dynamics in space (Bonte & Dahirel, 2017).

Phylogenetic information has been extensively used to infer dispersal distances for species without data (Barbet-Massin et al., 2012; Thomas, 2008). However, this approach neglects that dispersal can evolve rapidly by adaptive processes (Stevens et al., 2010), and that contrasting environmental conditions can generate variability in phenotypic dispersal patterns among individuals or populations (Bonte & Dahirel, 2017; Clobert et al., 2009). Our results show that both long and median dispersal distances have weak phylogenetic conservatism, indicating that population-level drivers such as landscape structure, or more labile behavioural traits, could play an essential role in determining dispersal (Blomberg et al., 2003; Nathan, 2001). Our results revealed lower phylogenetic signals in long-distance (compared to median) dispersal events, which could indicate that particularly long-distance movement are strongly context-dependent (Lowe, 2009). The overall phylogenetic lability on bird dispersal suggests that evolutionary history should only be used as predictor of dispersal ability when data are scarce and should otherwise be used with caution.

Accurately measuring age dispersal differences for many species has typically been hampered by the low juvenile survival rates compared to adults and because dispersal distances often exceed study area boundaries (Greenwood & Harvey, 1982; Newton, 1998). Here, we take advantage of continent-wide ringing and recovery efforts to show that natal dispersal of immature individuals that depart their natal range in search of new sites is generally more extensive and covers a wider geographical area than breeding dispersal (Greenwood & Harvey, 1982; Hollenbeck et al., 2018; Paradis et al., 1998). This considerable dispersal asymmetry between ages could arise from a range of selective pressures, such as inbreeding avoidance, competition among offspring, or simply finding suitable habitat (Clobert et al., 2012; Hendry et al., 2004). In contrast, mature breeders have evolved comparably lower breeding dispersal rates favouring territories they already know from previous breeding attempts (Kokko & Lundberg, 2001). Disentangling whether dispersal strategies are conditional on age is essential to understanding how demography and fitness influence the overall dispersal process (Bonte et al., 2012).

Studies of marked individuals are essential for understanding life histories and population dynamics. The EURING database provides an unrivalled source of mark-recapture information at a continental scale that is of immense value to ecology and conservation (Du Feu et al., 2016) and, as we have shown here, for estimating empirical dispersal distributions. However, sampling effort and detection in ring-recovery data vary considerably over time, space, species, and recovery types (Naef□Daenzer et al., 2017; Perdeck, 1977; Thorup et al., 2014). If not corrected for, this typically results in unsubstantiated estimates of dispersal that can lead to biased results or, in worst cases, wrong conclusions. Here, we identified sampling biases related to heterogeneous variation in ringer and finder activities (uneven spatial coverage, uneven sampling effort per type of recapture, heterogeneous reporting threshold between schemes) and biases related to the recoveries of birds on migration. We approached these biases by (1) using methods to exclude (filter) and standardise subsets of the data, keeping only the reliable observations (Geldmann et al., 2016) and (2) with an appropriate analytical approach to estimate dispersal for left-censored data using a Bayesian approach. This analysis and approach can be helpful for those working with large mark-recapture datasets from any taxa which cannot infer sampling effort or account for uneven detectability (using the provided code, see Data Accessibility). The filtering process and analysis could also be helpful to improve running monitoring programs or plan future ones.

The robust empirical characterisation of the avian dispersal kernels as presented in this study is crucial for conservation and management since and for predicting potential future range changes. The estimated dispersal distances as well as the analytical tools designed here provide many avenues for future research. Outstanding questions include, among others, the assessment of dispersal syndromes to understand how dispersal kernels vary across species traits and explore covariation patterns between dispersal and other traits (Clobert et al., 2009; Ronce & Clobert, 2012) and the exploration of how dispersal processes respond to habitat fragmentation and climate change (Bowler & Benton, 2005; Travis et al., 2013). The presented study paves the road towards a new generation of more realistic modelling and comparative studies to evaluate the role of dispersal in several issues of population biology and their eco-evolutionary dynamics under global change.

## Data and Code availability

Ring-recovery data is available upon request through the EURING Data Bank. Dispersal estimates and code will be available after publication from ZENODO repository: Guillermo Fandos (2021). guifandos/European_bird_dispersal: v0.1.0-Edispersal (v0.1.0_Edispersal). Zenodo. https://doi.org/10.5281/zenodo.5565077. Code available until publication https://github.com/UP-macroecology/European_bird_dispersal)

## Acknowledgements

We thank the Euring DataBank managers (most recently Dorian Moss) for curating and supplying the data, and the many thousands of ringers and members of the public who generated the data in the first place. Special thanks to Stephen R. Baillie for helpful comments on a previous version of this paper.

## Author contributions

**Guillermo Fandos**: Conceptualization (lead); Investigation (equal); Data request (lead); Formal analysis (lead); Methodology (lead); Writing – original draft (lead); Writing-review & editing (equal). **Matthew Talluto**: Conceptualization (supporting); Formal analysis (equal); Methodology (equal); Writing – original draft (supporting). Writing-review & editing (equal). **Wolfgang Fiedler:** Conceptualization (supporting); Investigation (supporting); Data request (supporting); Writing – original draft (supporting); Writing-review & editing (equal). **Robert A. Robinson**: Conceptualization (supporting); Investigation (supporting); Data request (supporting); Methodology (supporting); Writing – original draft (supporting); Writing-review & editing (equal). **Kasper Thorup:** Conceptualization (supporting); Investigation (supporting); Writing – original draft (supporting); Writing-review & editing (equal). **Damaris Zurell**: Conceptualization (equal); Funding acquisition (lead); Investigation (supporting); Data request (equal); Formal analysis (supporting); Methodology (supporting); Writing – original draft (equal), Writing-review & editing (equal).

