## supplemental material for "Standardised empirical dispersal kernels emphasise the pervasiveness of long-distance dispersal in European birds"

### **Supplemental Analysis**

S1. Details about the data selection process

S2. Dispersal estimates from different types of recovery

S3. Birds on migration bias

S4. Fitting dispersal kernel in a Bayesian framework

S5. Natal and breeding dispersal distance relationship with the Weibull distribution

S6. Relationship between Paradis et al. 1998 mean dispersal distances and those estimated here for 75 species

S7. Long-distance dispersal (km) along the bird phylogeny for the average (234 species), breeding (113 species) and natal dispersal (121 species).

S8. Distribution of the estimated median dispersal distance (km) for all species (n=234) and for species with enough data to separately calculate breeding and natal dispersal (n=1113).

S9. For each species is indicated the number of ring recoveries used for the analysis (n), the function used (function_id; Exponential, Gamma, Weibull, Half-Cauchy), the posterior model probabilities from marginal likelihoods (function_comparison), the age type of dispersal (type: average, breeding or natal dispersal), and the parameter mean (mean), standard error (se_mean), standard deviation (sd), 95% credible interval (2.50%-97.50%). Gamma and Weibull functions have a column for each one (parameter1, parameter2). Exponential and Half-Cauchy have NA in parameter 2.

S10. For each species is indicated the median dispersal distance (median), 95% credible interval (2.50%; low_CI-97.50%; upper_CI), the lower-bound 5% of the posterior predictive distribution (lower_distance), the upper-bound 95% posterior predictive distribution (upper_distance/ long-distance dispersal), the number of ring recoveries used for the analysis (n), the distribution function used (function_id; Exponential, Gamma, Weibull, Half-Cauchy), the age type of dispersal (type: average, breeding or natal dispersal), and the posterior model probabilities from marginal likelihoods. 95% of individual birds will disperse a distance within the interval between distance and upper_distance.

#### S1. Data selection

1. Schemes excluded from the analysis Israel, Cyprus (Nicosia), Cyprus (Kuskor), Malta and Russian Federation.
2. Species data excluded from 1) Norway (*Haliaeetus albicilla, Accipiter gentilis, Aquila chrysaetos, Pandion haliaetus, Falco peregrinus, Gallinago media, Bubo bubo, Strix uralensis, Strix nebulosi*). 2) Hungary: for ten selected species (*Riparia riparia, Ichthyaetus melanocephalus, Ciconia ciconia, Coracias garrulus, Platalea leucorodia, Chroicocephalus ridibundus, Falco vespertinus, Ardea alba, Cygnus olor, Ciconia nigra),* only birds that were recaptured outside Hungary were included, not within Hungary.
3. In order to achieve the 60% dead and 40% alive recoveries in the data request to test the effect on the dispersal estimates of the recovery type, we apply the next filters:

- If the total number of records in a kernel (ignoring dead/alive) is under 20, delete them.
- If the total number of records in a kernel (ignoring dead/alive) is between 20 and 100, keep them.
- If the total number of records in a grid (ignoring dead/alive) is over 100, including at least 60 dead and 40 alive, we randomly select 60 dead and 40 alive (but then delete any orphan records so that the total number will be between 98 and 100).
- If there are more than 100 records but less than 60 dead or less than 40 alive. Complete without considering the recovery type

#### S2. Dispersal estimates from different types of recovery

EURING database observations of ringed birds had a field code for the condition of the bird when it was found (Du Feu et al., 2016). We grouped this condition in three major ways: first, live recapture at subsequent sampling occasions; second, resighting of tagged birds without physical capture; and third, ring recovery from birds that are found dead, commonly as a result of hunting.

The spatial distributions of birds encountered in these three different conditions are likely to be heterogeneous in space and time. Therefore, we used different approaches to test for potential differences in dispersal distances estimated from the different recovered condition.

First, Paradis et al (1998) argue to only work with dead recoveries since live captures have a strong bias where field ornithologist tends to catch birds. To test this hypothesis, we develop a preliminary analysis where we extracted the Human Foot Print values (HFP; Venter et al., 2016) in all the ring-recoveries using the extract function of the raster R package (Hijmans & Van Etten, 2016). Then, we used General linear mixed models (GLMM to examine whether the probability of finding a dead bird and the probability of a live recapture have a bias towards areas with high human activity. We used species as random effects and type of recovery as a fixed effect. HFP values were first standardised to a mean of 0 and SD of 1. The GLMM was performed using lme4 package (Bates et al., 2015). The results show no significant differences between type of recoveries (Figure S1).

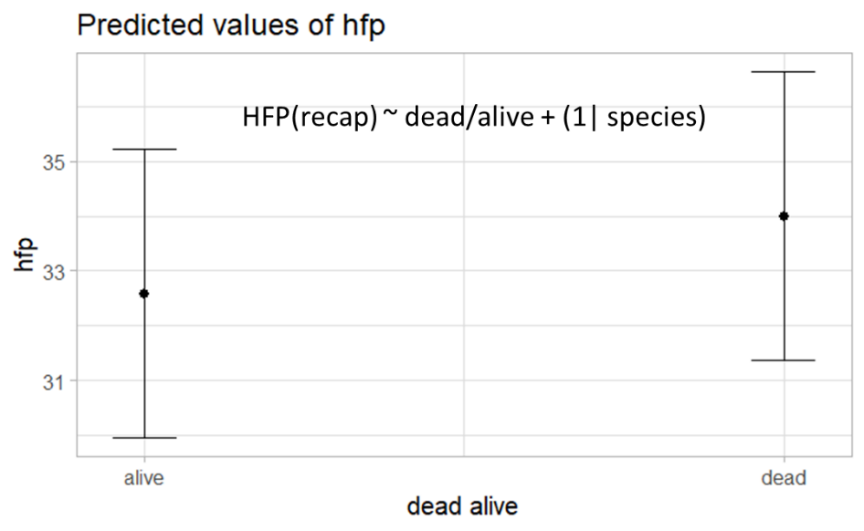

*Figure S2.1: Average (±95% C.I.) Human Foot Print (hfp) values of the sites where the recoveries were found according to the recovery type (dead or alive).*

Second, we explore and compare the dispersal distances estimated from the different recovered types (dead, recapture, and resights). To avoid problems related to estimating dispersal for species with a low number of recoveries, we restricted the analysis to bird species with>50 records for the three types of condition. A total of 42 species were included in the analyses.

We estimated dispersal kernels for the three types of recoveries and compared the dispersal median estimates between them using General Linear Models (GLM). We used as a response variable the dispersal median distance from dead recoveries and the dispersal median distance from alive recoveries, and the type of alive recovery, recapture or resights, as fixed effects.

*Table S2.1: Linear mixed-effects models describing the differences dispersal distances estimated from dead recaptures with the dispersal distances estimated from alive recaptures (Model Dead/Recapture) and from alive resights (Model Dead/Resight). The standard errors are shown in brackets, and the p-values are indicated as * in the table. *** p < 0.001; ** p < 0.01; * p < 0.05.*

|  | Model Dead/Recapture | Model Dead/Resight |
| --- | --- | --- |
| (Intercept) | 15.04 *** | 18.98 ** |
|  | (3.28) | (5.48) |
| dispersal_recapture | 0.63 *** |  |
|  | (0.07) |  |
| dispersal_resight |  | 0.43 ** |
|  |  | (0.14) |
| N | 42 | 42 |
| R2 | 0.67 | 0.20 |

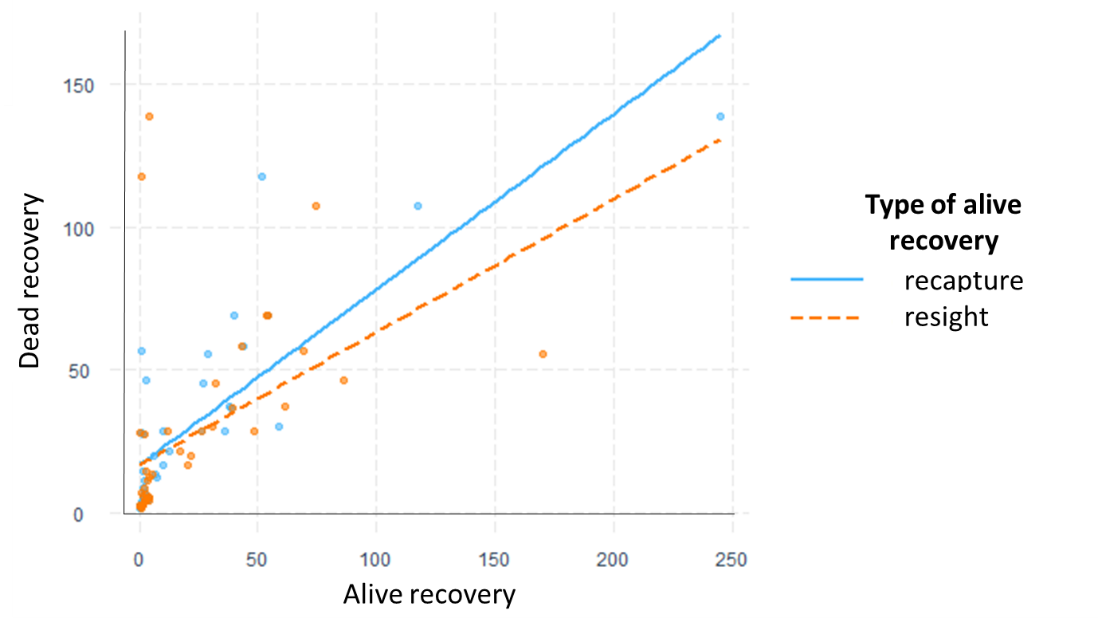

*Figure S2.2: Correlation between dispersal distance estimated from dead and alive recoveries according to the type of alive recovery (recapture: blue, resight: orange).*

The results show that although dispersal estimates from both recapture and resight recoveries were positively correlated with dead recoveries’ estimates (Figure S2). The explanatory variance drastically reduces from 67% to 20% when we used resight instead of recaptures (Table S1). Thus, we can conclude that although correlated, the process behind the catching effort by ringers and reporting probability between recapture and resight recoveries can impact the dispersal estimates. This difference is probably a consequence of the nature of the resight recoveries from specific projects or species. Therefore, we decided to exclude the resights from the subsequent analysis .

#### S3. Birds on migration bias

This section aims to identify captures/recaptures of migrating individuals and exclude them from subsequent analyses. Falsely including them could lead to an overestimation of dispersal distances (Paradis et al., 1998).

**Preliminary steps to prepare the data for the analysis:**

- Join EURING database with species traits. We use the trait list from Storchová L & Hořák D (2018). However, seven species have not traits in the list, so we used close related species to fill those ones. (*Luscinia svecica, Hippolais pallida opacus, Corvus corone cornix , Carduelis flavirostris, Carduelis flammea, Carduelis flammea cabaret, Miliaria calandra*). We delete the Caribean flamingo (*Phoenicopterus ruber*) from the EURING database
- Scale distance by the elapsed time between capture and recapture. We separate breeding and natal dispersal.
  - For breeding dispersal: Individuals with more time (years) between capture and recapture probably have larger dispersal distances by cumulative dispersal events. To scale all the individuals, we assume that all the individuals disperse all the years, and all the individuals disperse constant distances. We apply a scale by a random walk: *dispersal_scaled= distance/sqrt(Number of years between capture and recapture)*
  - For natal dispersal: We can identify three groups of natal dispersal. 1- Capture as juvenile and recapture as an adult in the first year of reproduction*. Not scaled*. 2- Capture as juvenile and recapture as an adult after the first year of reproduction. Scale the years after the first year of reproduction.3- Capture as juvenile and recapture as an adult before the first year of reproduction. *Not scaled*
- Exclude recaptures below latitude 35 degrees and longitude larger than 40 degrees. This step will exclude some migratory individuals that were wintering in Africa, for example.

**Exclusion of migratory individual’s analysis**

For identifying migrating individuals, we apply the following steps to only the migratory species (Storchová & Hořák, 2018):

1- **Select a potential breeding period for each species and grid cell in Europe.**

First, we will estimate the breeding core period, and from there, calculate a threshold distance where we can consider that an individual could be during a migratory movement. Potentially, the breeding core period has high spatial and temporal variability across Europe (Gordo, 2007). Differences in the start of breeding between North and South Europe could be of more than one month for the same species. To estimate this “breeding core period”, we will use the following steps. We analyse the distance between capture and recapture against time, assuming that the drop in distance at the beginning of the period could be related to migratory individuals’ arrival. In this first step, our goal is to find a period (“breeding core period”) without significant changes in the distance between the capture and recapture to later exclude potentially migratory movements from the dispersal analysis.

- Select only migratory species (at least one population has to be partial migrant, long-distance migrant, or short-distance migrant).
- Associate each recapture with a grid cell (5°) in Europe (Figure S4B).
- We select specific breeding periods (run GAM models) for each species and grid cell using a moving window approach. In this moving window, we select the data for each grid cell, and we add to the analysis the eight adjacent (5°) grid cells to increase the amount of data for the GAM analysis. We exclude the specific grid cell from the analysis when the complete data selection (the grid cell selected + 8 adjacent cells) has less than 20 records.
- We run the GAM model, with the distance scaled between capture and recapture against time as a smoothing factor. We used a Negative Binomial GAM to model it with a log link function. We set ten dimensions as the upper limit on the degrees of freedom associated with the smooth term (the exact value is not critical, but it is essential not to make it restrictively small or large and not too computationally costly; Wood, 2017)
- We developed a sensitivity analysis to select the best term (k) of effective degrees of freedom to detect a breeding core period. We test four different k terms (5,10,15,20). Finally, we selected that a smooth term of 10 was allowing us to select a proper breeding period for a large number of species and grids.
- To identify periods of significant change in the distance between the capture and recapture in the breeding period, we assess the slope by computing the first derivative of the GAM spline and the confidence intervals from their standard errors. We evaluated the derivative and looked for locations where the confidence interval does not include zero, considering them the flat plateau.
- When it is longer than 20 consecutive days, we can consider this plateau (not significantly increasing or decreasing the distance between capture and recapture) as our breeding core period. If there is more than one period > 20 days, we select the largest one (see Figure S4D).

BOX S3: Spatiotemporal GAM analysis

a) We developed a preliminary analysis where we developed spatiotemporal GAM models for some species (*Ciconia ciconia, Phalacrocorax carbo, Milvus migrans, Falco Tinnunculus*). Here, we tested if the distance between capture and recapture varies across Europe and changed over the breeding period. For that, we used a tensor product spline to test if there is a significant interaction between space (latitude and longitude) and time (days of the breeding period), explaining the distance between the capture and recapture. In addition, we account for local heterogeneity with the scheme that reports each recapture as a random effect smooth. The results show some specific spatiotemporal trends of the distance between the capture and the recapture. We predict these previous models in each grid cell. However, the first derivative from these predictions was not successful in finding a proper breeding period.

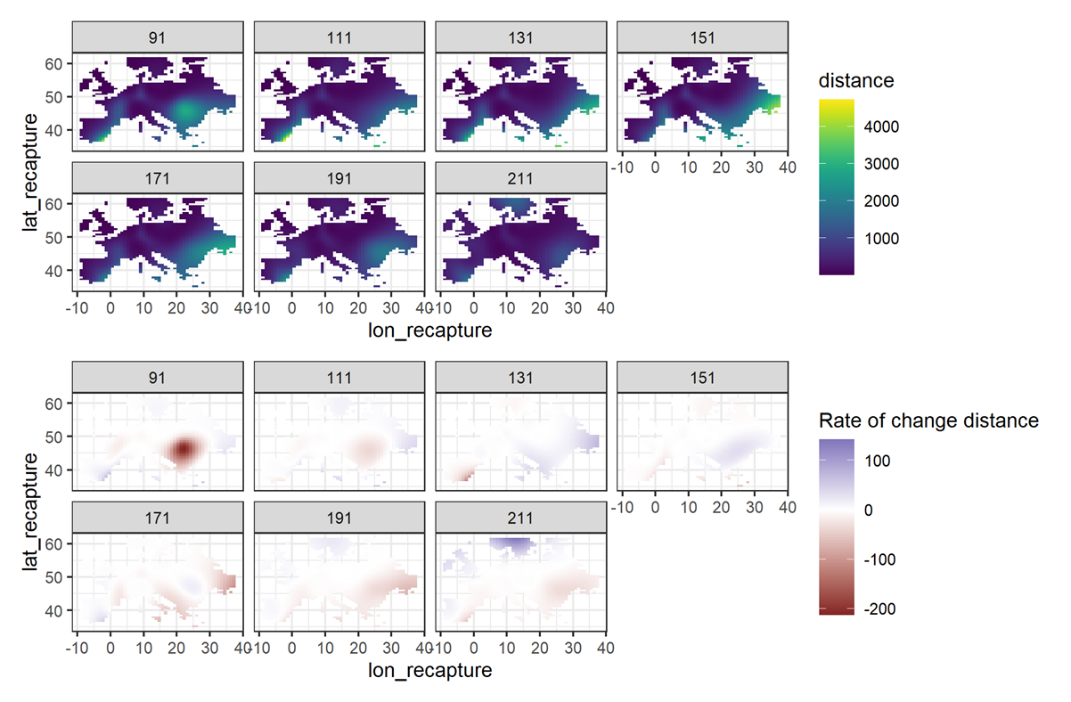

Figure S3.1: Example with the white stork (*Ciconia ciconia*). Top panels are the prediction from the GAM models with the spatio-temporal trends in the distance between the capture and recapture. Down panels show the rate of change at each location and at each time point (every 20 days) and determine which locations are changing most quickly.

2- **Threshold distance to detect migratory individuals**

- We use the scale distances observed in the core-breeding period as a cut-off distance for filtering all other re-encounters. Thereby, we define the 90/95 % quantile (our idea is to run the dispersal kernels with both thresholds) of the core-breeding breeding period distances (specific for each grid and each species) as the cut-off distance.
- We discarded any individuals exhibiting distances larger than this cut-off distance (Figure S4A)
- We visual inspect if we were successful in excluding potential migratory individuals by comparing the raw data with the threshold of 90% quantile and 95% quantile

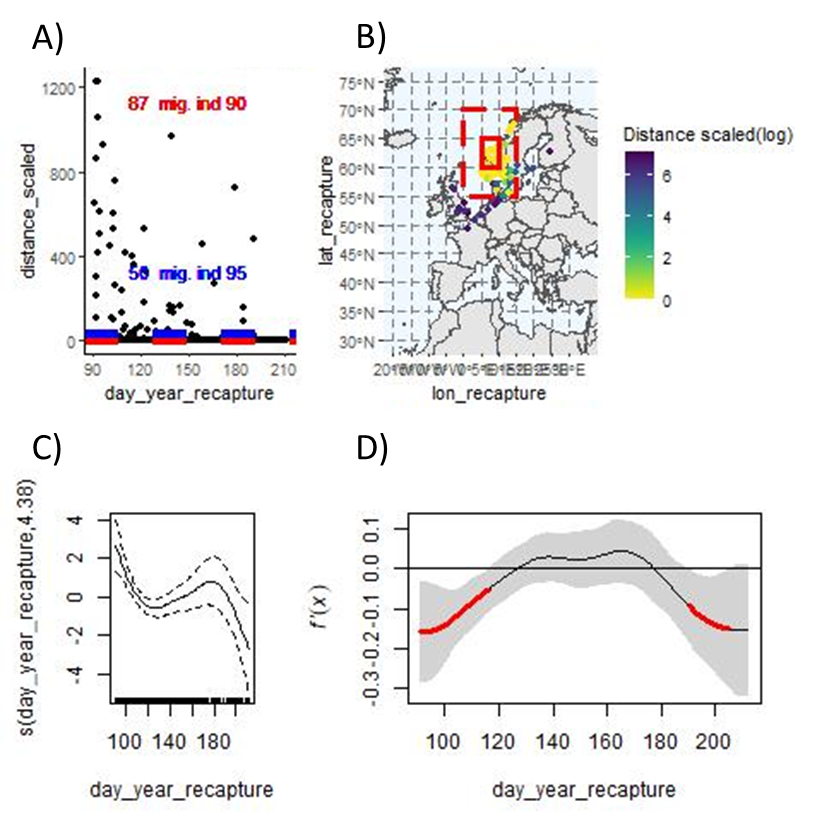

Figure S3.2: a) Scatterplot between the scale distance and the day of the year the birds were recaptured for *Fringilla coelebs*. The blue dash line indicates the threshold for the 95% quantile and the red dash line for the 90% quantile, both with the respective number of potential migratory individuals excluded with each threshold b) Map with the grid 60 latitude/10 longitude (solid red line) and the eight adjacent cells (dashed red line) for the *Fringilla coelebs* with the data used for the analysis colour coded with the log distance between the capture and recapture. c) GAM analysis with the distance scaled between capture and recapture against time as smoothing factor with the confidence intervals 95%. d) The first derivative of the GAM spline, in red, is the part that where the trend is significantly decreasing (95% confidence interval). Hence, in this period, we considered the breeding core period to calculate the 90 % and 95 % quantile of the distances from day 119 until 185 for *Fringilla coelebs* in this specific grid cell.

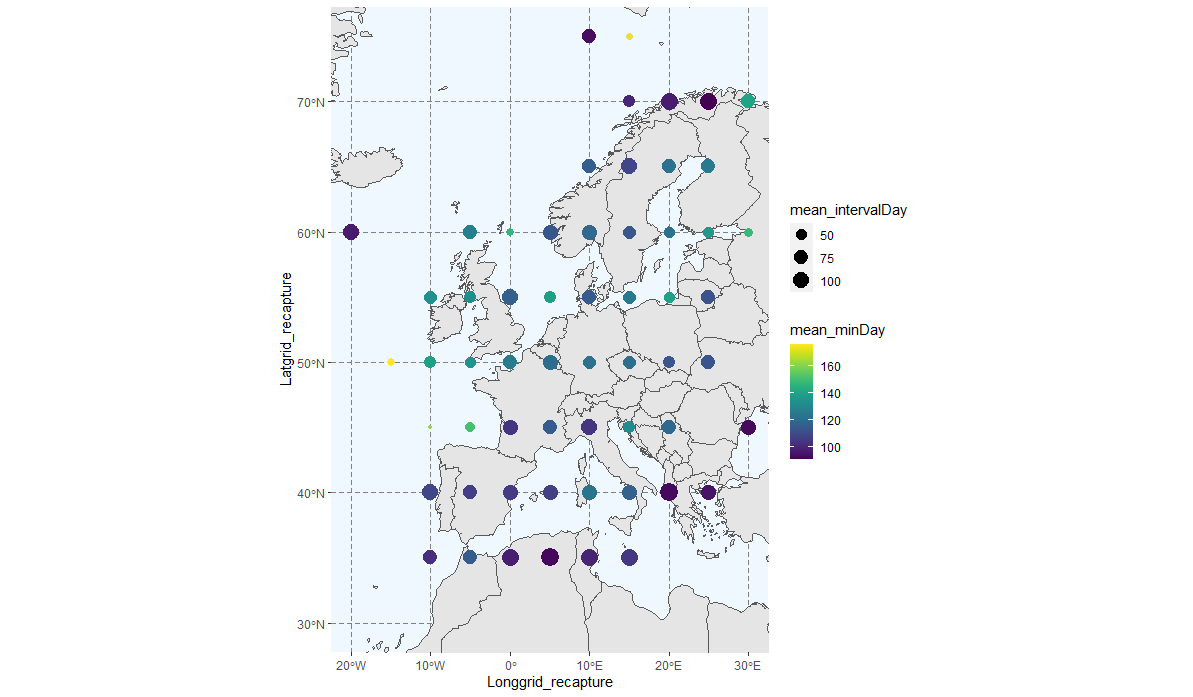

Figure S3.3: Average between all species of the minimum day (point colour; yellow later start of the breeding period) and the number of days during the breeding period core (interval days; the size of the point) estimated across all Europe.

The preliminary visualisation with the raw data shows that some individuals might be during migratory movements, as the largest dispersal distances correspond to migratory directions (Northeast/Southwest). These large distances in those specific directions were mostly discarded after the GAM process and better with the 90% threshold (Figure S6-7).

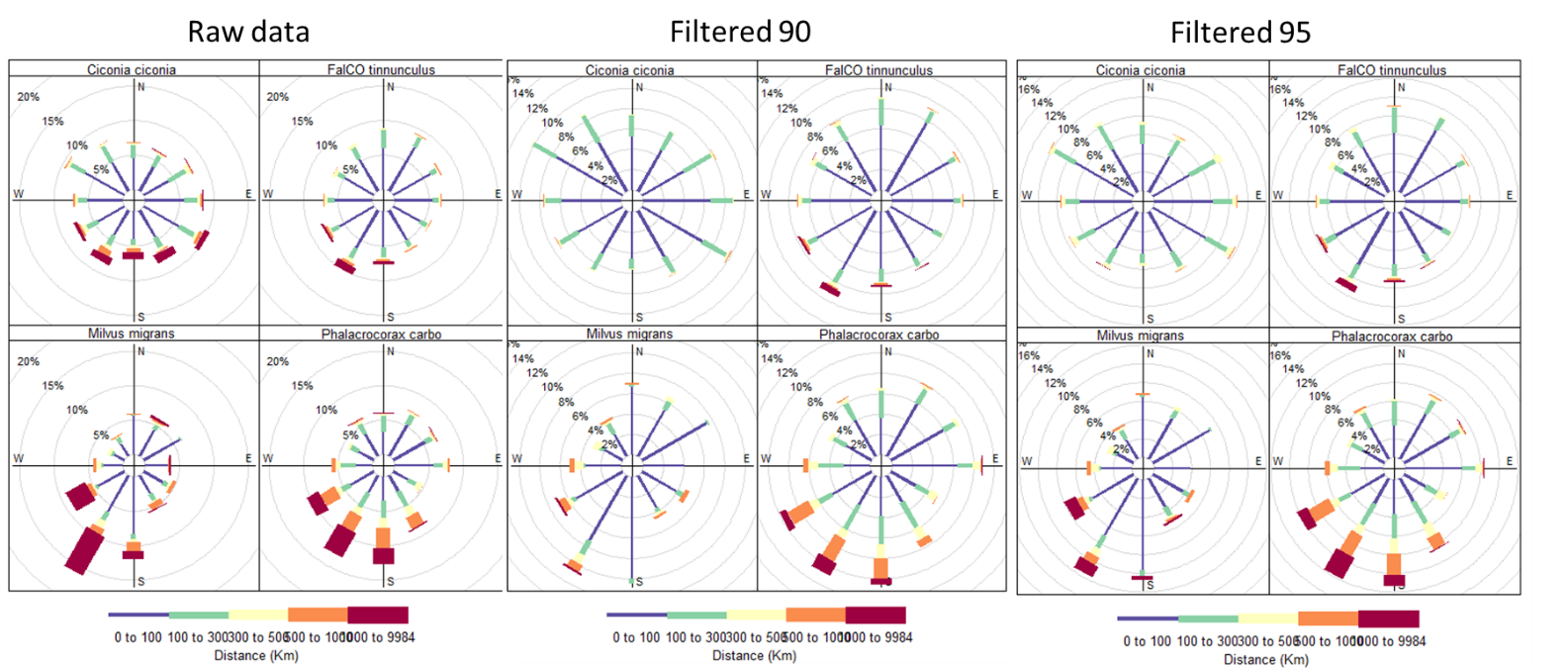

Figure S3.4. Distance and directions between ringing and recapture for some example species *(“Phalacrocorax carbo”, "Falco tinnunculus", "Ciconia ciconia", "Milvus migrans").* The plots summarise data from all available ringing schemes and show, for each species, the proportion of individuals going in each direction (360°), with the colour representing the distance between the capture and the recapture (red colours larger distances than blue colours) for before (raw data) and after applying the GAM analysis with 90% threshold (Filtered 90) and 95% threshold (Filtered 95) to exclude potential migratory individuals.

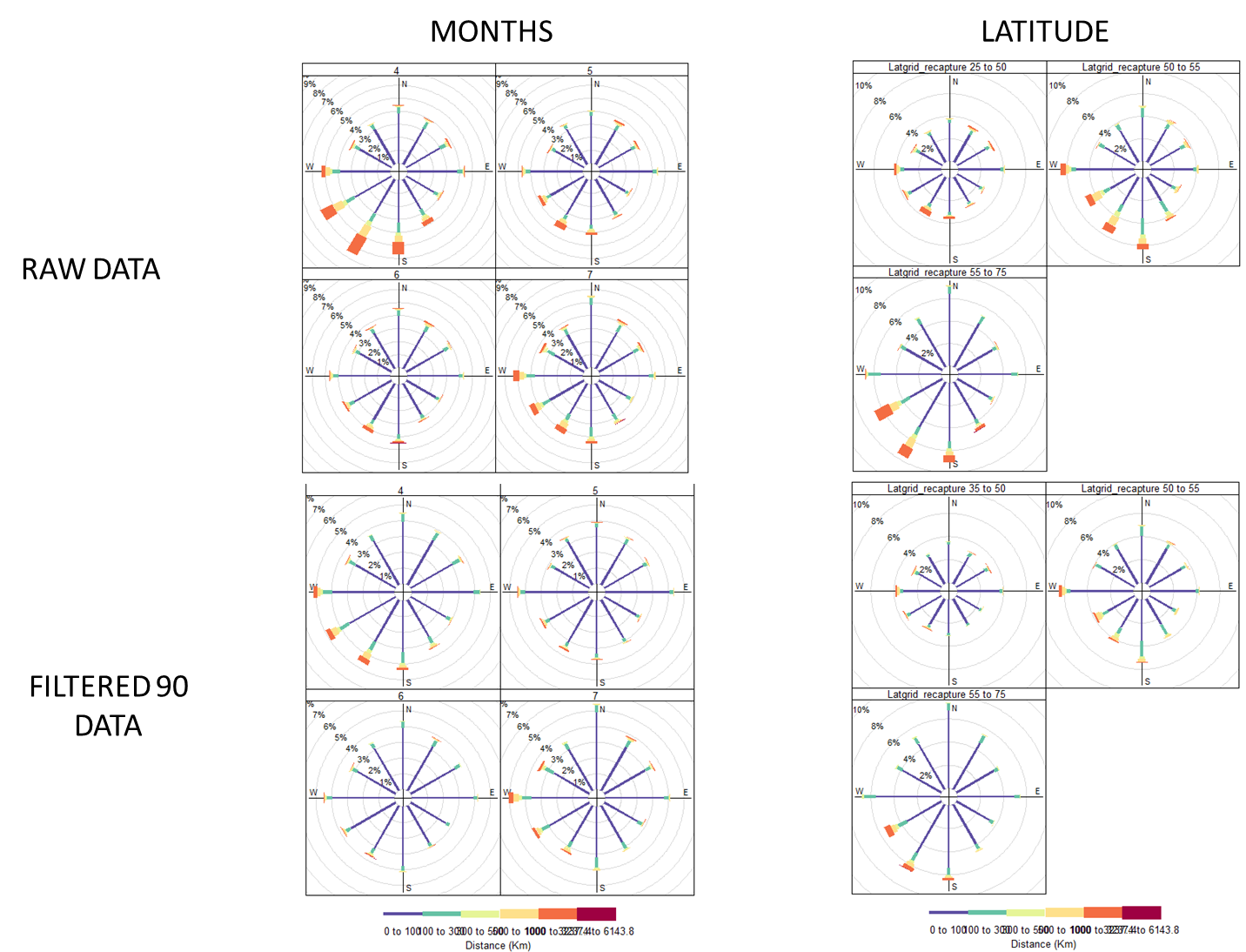

Figure S3.5: Distance and directions across months of recapture (left panels) and latitude (right panels) for all the species before (raw data; top panels) and after applying the GAM analysis with the threshold of 90% to exclude potential migratory individuals (down panels).

#### S4. Fitting dispersal kernel in a Bayesian framework

Prior distribution for the threshold parameter

Based on exploratory analyses and scheme reports (EURING web), we used a gamma prior distribution with shape = 2500, and scale = 500 to estimate the threshold parameter for each scheme.

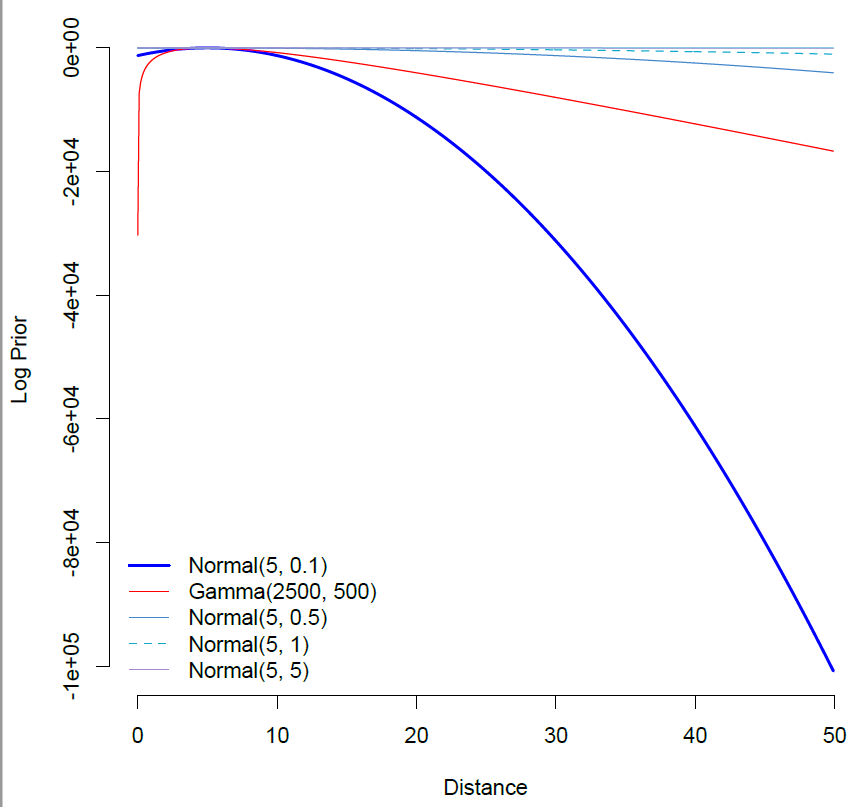

Figure S4.1: Different prior distribution explored to estimate the threshold parameter for each scheme.

#### S5. Natal and breeding dispersal distance relationship with the Weibull distribution

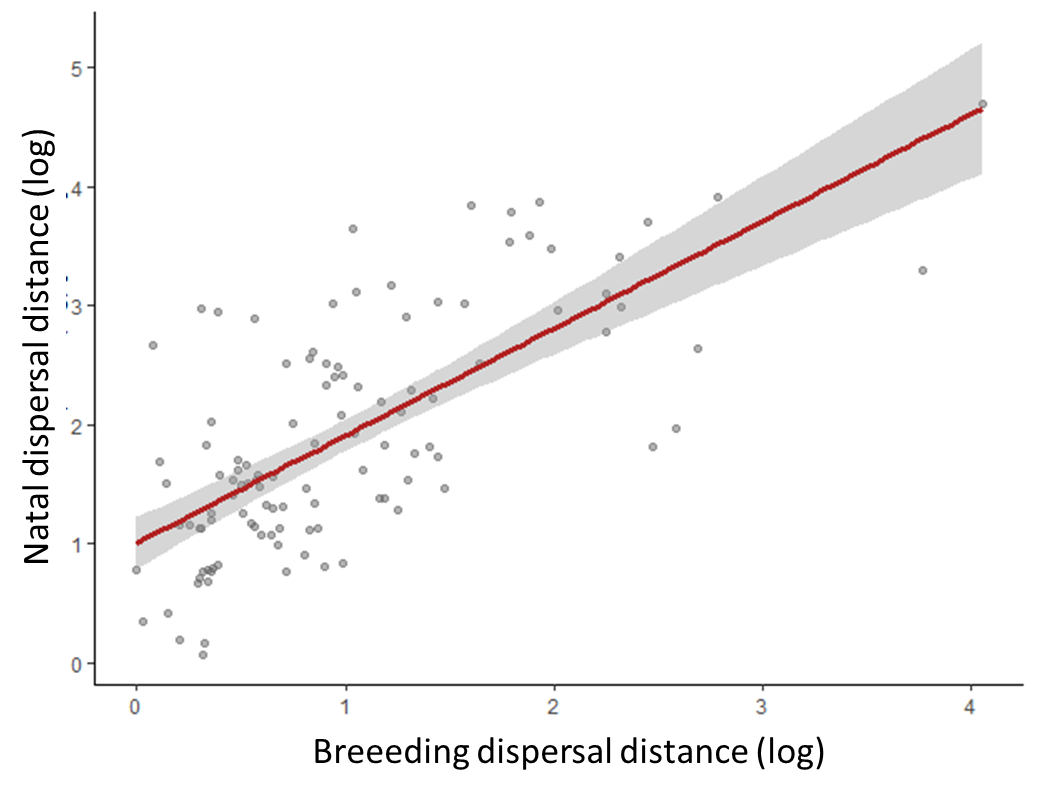

Figure S5.1: Linear relationship between breeding and natal dispersal (log) both estimated with the Weibull distribution. The multivariate generalised linear mixed model results with the natal dispersal distance (log) estimate as a response variable, the breeding dispersal distance (log) as a fixed effect, and phylogeny as a random effect are post.mean= 0.866; 95% CI= 0.677-1.068; pMCMC < 0.001

#### S6. Relationship between Paradis et al. 1998 mean dispersal distances and those estimated here for 75 species

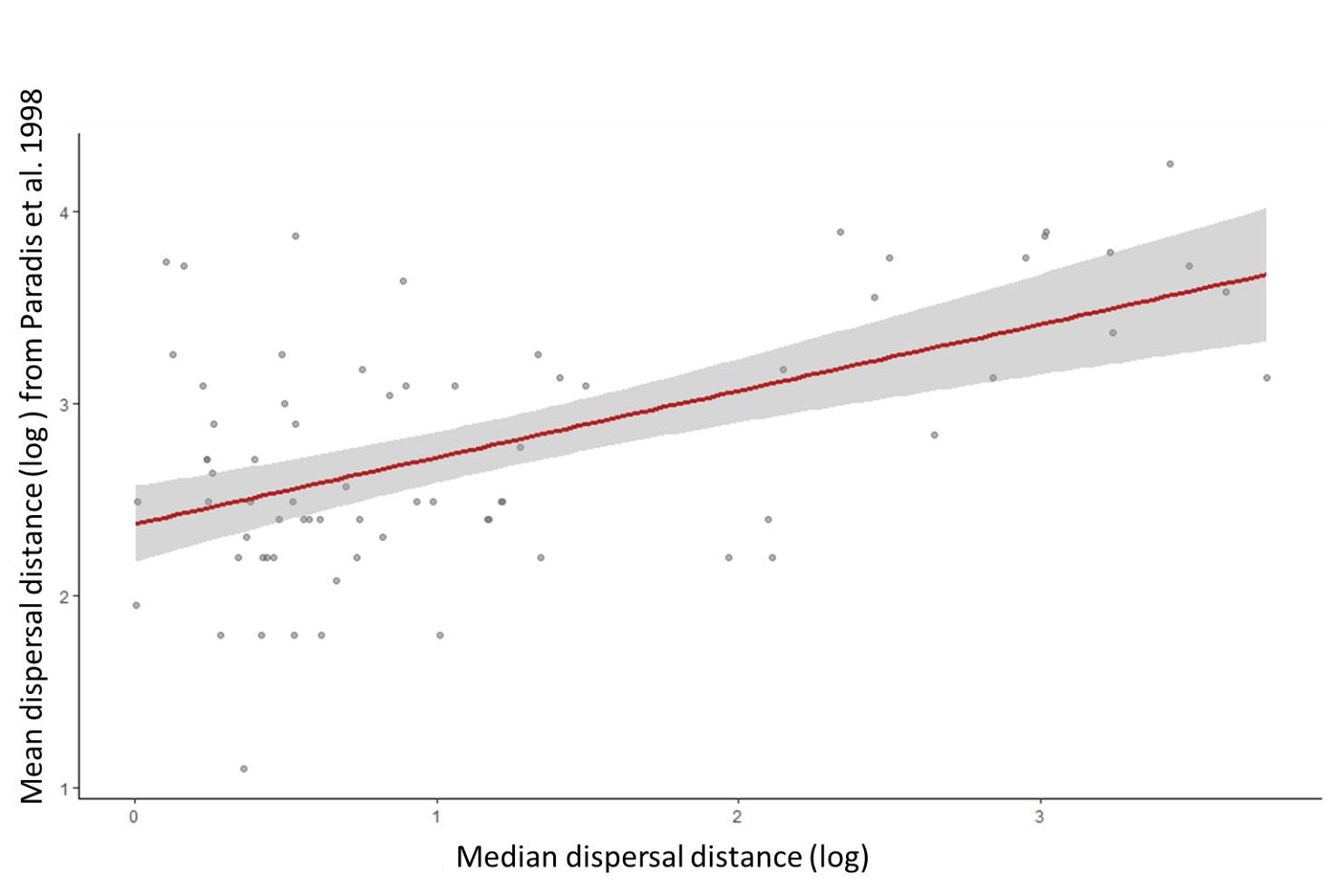

Figure S6.1: Linear relationship between mean dispersal (log) estimated in Paradis et al. 1998 and the median average dispersal distance (log) estimated here. The multivariate generalised linear mixed model results with the Paradis et al 1998 dispersal mean estimate as a response variable, the median breeding dispersal distance as a fixed effect, and phylogeny as a random effect are post.mean= 0.366; 95% CI= 0.211-0.524; pMCMC < 0.001

#### S7. Long-distance dispersal (km) along the bird phylogeny for the lifetime (234 species), breeding (113 species) and natal dispersal (121 species).

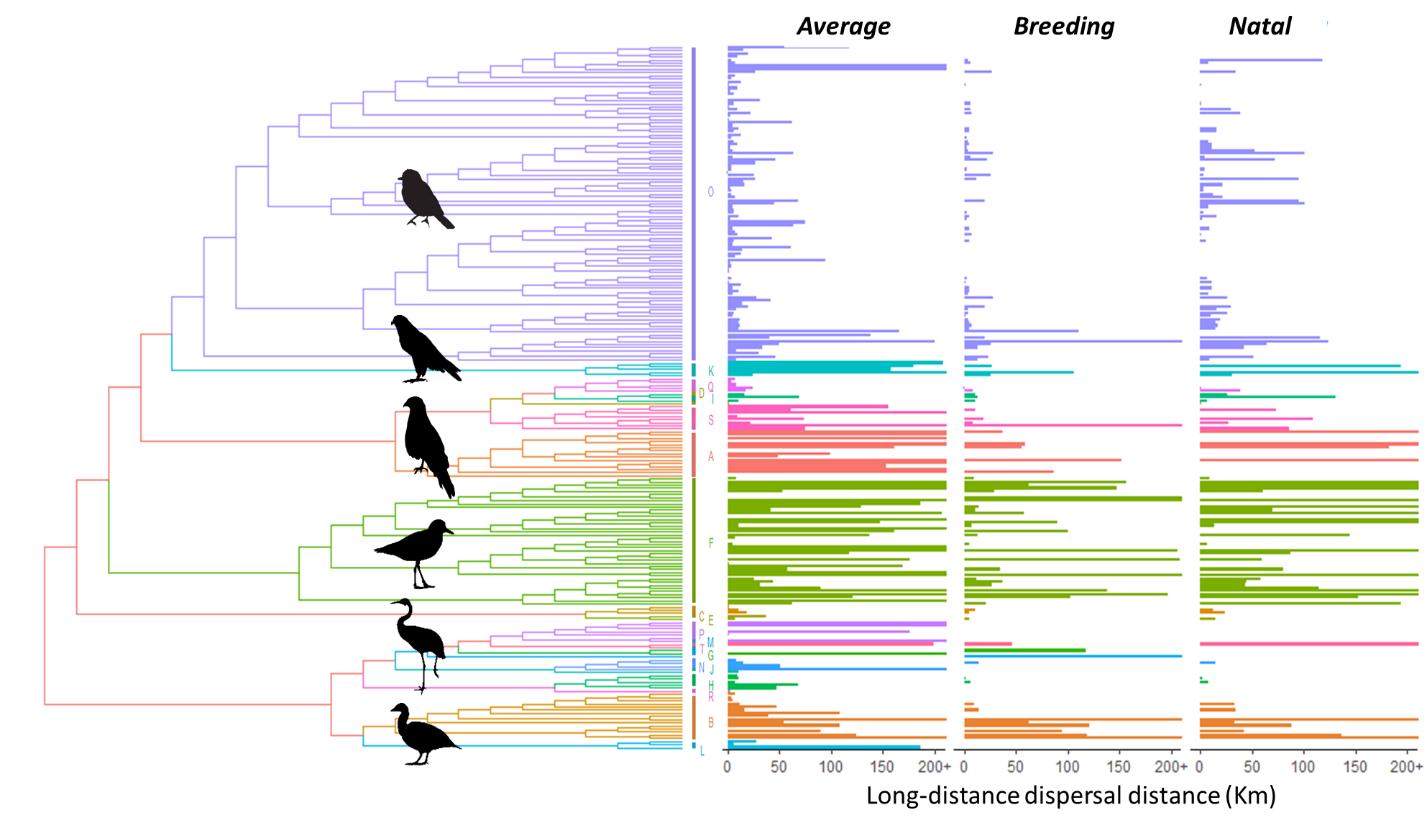

Figure S7.1: Long dispersal distance (km) along the bird phylogeny for the average (234 species), breeding (113 species) and natal dispersal (121 species). The dispersal distance is truncated at 50 Km for visualisation purposes. Each colour and letter represent the same Order in the phylogeny and the bar plots. *A: Accipitiformes, B: Anseriformes, C: Apodiformes, D: Bucerotiformes, E: Caprimulgiformes, F: Charadriiformes, G: Ciconiiformes, H: Columbiformes, I: Coraciiformes, J: Cuculiformes, K: Falconiformes, L: Galliformes, M: Gaviformes, N: Gruiformes; O: Passeriformes, P: Pelecaniformes, Q: Piciformes, R: Podicipediformes, S: Strigiformes, T: Suliformes.*

#### S8. Distribution of the estimated median dispersal distance (km) for all species and for species with enough data to calculate breeding and natal dispersal.

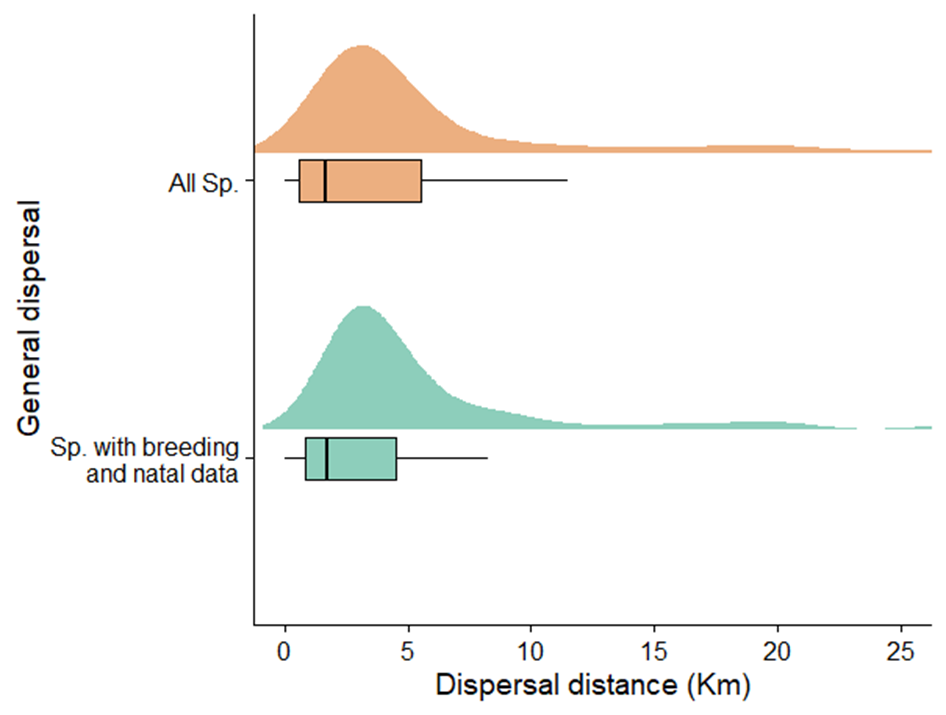

Figure S8.1: Distribution of the estimated median dispersal distance (km) for all species (n=234) and for species with enough data to separately calculate breeding and natal dispersal (n=113).

#### S9. For each species is indicated the number of ring recoveries used for the analysis (n), the function used (function_id; Exponential, Gamma, Weibull, Half-Cauchy), the posterior model probabilities from marginal likelihoods (function_comparison), the age type of dispersal (type: average, breeding or natal dispersal), and the parameter mean (mean), standard error (se_mean), standard deviation (sd), 95% credible interval (2.50%-97.50%). Gamma and Weibull functions have a column for each one (parameter1, parameter2). Exponential and Half-Cauchy have NA in parameter 2.

Table S9: Table_S9_species_dispersal_parameters.csv

#### S10. For each species is indicated the median dispersal distance (median), 95% credible interval (2.50%; low_CI-97.50%; upper_CI), the lower-bound 5% of the posterior predictive distribution (lower_distance), the upper-bound 95% posterior predictive distribution (upper_distance/ long-distance dispersal), the number of ring recoveries used for the analysis (n), the distribution function used (function_id; Exponential, Gamma, Weibull, Half-Cauchy), the age type of dispersal (type: average, breeding or natal dispersal), and the posterior model probabilities from marginal likelihoods. 95% of individual birds will disperse a distance within the interval between distance and upper_distance.

Table_S10: Table_S10_ species_dispersal_distances.csv
